# Maresin-2 Fine-tunes ULK1 O-GlcNAcylation to Improve Post Myocardial Infarction Remodeling

**DOI:** 10.1101/2023.07.16.549182

**Authors:** Jingjing Zhang, Chenyu Li, Yanzhao Wei, Shujuan Jiang, Xiaolin Wu, Qing Zhou, Shuang Yang, He Hu, He Huang, Bin Kong, Wei Shuai

## Abstract

Maresin-2, a specialized pro-solving mediator of inflammation has been consolidated to be a novel cytokine fine-tuning inflammatory cascade. However, the underlying molecular basis is still largely unknown. Focused on cardiac dysfunction and remodeling, we employed in vivo- and in vitro- based genome editing methodology tools including adenosine associated virus, adenosine virus, lenti-virus, plasmid transfection, and CRISPR-Cas9 methodology for investigation. As suggested, exogenous maresin-2 supplement facilitated autophagosome formation by microtubule-associated proteins 1A/1B light chain 3B (LC3) conjugation system under the modulation of O-GlcNAcylation dependent ULK1 activation, whereas reversed by ULK1 S409A and S422A mutagenesis, showcasing the potential O-GlcNAc (O-linked β- N-acetylglucosamine) modifiable sites on ULK1. Moreover, we found that hereafter maresin-2 treatment glutamine-fructose-6-phosphate aminotransferase 1 (GFAT1) which is accessary to sense hexosamine biosynthesis influx is more likely the prime checkpoint for conjugating O-terminal β-N-acetylglucosamine motif onto ULK1, rather than O-linked N-acetylglucosaminyltransferase (OGT). Mechanistically, maresin-2 largely prohibits transforming growth factor-β (TGF-β)-activated kinase 1 (TAK1), therefore increasing the availability of TAB1 for GFAT1, which encourages O-GlcNAcylation of ULK1.

## Introduction

Myocardial infarction with a pathogenic basis of thrombosis or vascular occlusion remains a global medical concern overhauling characterized by high morbidity and mortality^1^. Post infarction myocardial repair is compounded by complexities of stress response reactions associated with initiation and resolution of inflammation^2^.

Resolution of inflammation is a process in part mediated by specialized prosolving mediators (SPMs) to antagonize against inflammatory cascades^3^. Defective metabolism of SPMs and relevant analogs are common comorbidity of inflammatory-related diseases^3, 4^. As a subdivision, maresin-2 is biosynthesized from a long-chain n-3 polyunsaturated fatty acid—docosahexaenoic acid (DHA) mediated by 12-lipoxygenase (12-LOX)^5^. That 14-hydroperoxide intermediate catabolized from DHA by 12-LOX is the substrate for enzymatic epoxidation into the 13S, 14S-epoxy- maresin. Manifested by divergent hydrolysis entrance, 13S, 14S-epoxy maresin is catabolized into maresin-1 (via enzymatic hydrolysis) or maresin-2 (via soluble epoxide hydrolase)^6, 7^. As the biological actions of maresin-1 have been extensively documented, the role of maresin-2 is largely unknown.

Macro-autophagy (hereafter autophagy) is an immediate cellular degrading mechanism pivotal in assembly of autophagosome in which comprised of organelles and cellular debris, be they cytoplasm or membrane localized, are sequestered by cargo structure into lysosome in a fusion manner for bulk recycling of cellular components^8^. In mammalian autophagy, the Unc-51–like autophagy activating kinase (ULK1) complex is a core autophagy initiating machinery that connects cargo recognition with isolation membrane biogenesis and elongation of nucleated autophagosome^9,10^. There are four proteins to form ULK1 complex: ULK1 kinase, autophagy related 13 (ATG13), autophagy related 101 (ATG101), and focal adhesion kinase family interacting protein of 200 kDa (FIP200)^11∼13^. ULK1 complex selectively recruits phospholipid phosphatidylinositol 3-phosphate (PI3P) module via interplay with VPS34 (vacuolar protein sorting 34) complex. These complexes and proteins, as a result pack early phase autophagosome, such that underpins autophagic flux comprised of debris cargo identification, autophagosome closure and fusion with lysosomes which is implicative by catalyzing the conjugation of the ubiquitin-like LC3 from LC3-I to LC3-II^14∼17^.

O-GlcNAcylation, a posttranscriptional modification process of attaching O-linked β-N-acetylglucosamine (O-GlcNAc) moieties on serine/threonine residues is regulated by O-GlcNAc transferase (OGT) catalyzing the incorporation of β-O-GlcNAc from the donor substrate uridine-5′-diphospho-*N*-acetylglucosamine (UDP-GlcNAc) and O-GlcNAcase (OGA) catalyzing the removal of O-GlcNAc from the glycosylated substrates^18, 19^. Glucose, glutamine and glucotamine are substrates for hexosamine biosynthetic pathway (HBP) for deriving of UDP-GlcNAc^20^. Baseline HBP routine utilizes 2∼5% total glucose and mainly modulated by glucose/glutamine supply amount and the rate-limiting enzyme glutamine: fructose-6-phosphate amidotransferases (GFATs)^21, 22^. GFATs that converts fructose-6-phosphate and glutamine to glucosamine-6-phosphate (GlcN-6-P), the precursor of UDP-GlcNAc^23, 24^. Aberrant HBP in heart has been implicated to orchestrate cardiac cell fate from cues of glucose, fatty acid, and amino acid nutritious property^25^. Furthermore, cardiac GFAT activity with putative function on nucleus factor pathway might be regulated by inflammation cascade. That nuclear factor kappa-light-chain-enhancer of activated B cells (NF-κB) signaling interacting with GFAT1 in a fine-tuned manner modulates recruitment of downstream p38 mitogen-activated protein kinase (p38 MAPK) to promote autophagy occurrence^26^. If maresin-2 could be involved in cardiac repair and remodeling, elucidating its underlying mechanism including autophagy and O-GlcNAcylation, is helpful to provide theoretical rationality of maresin-2 for myocardial treatment.

Collectively, our findings suggest a previously unrecognized role of maresin-2 in cardiac homeostasis. Mechanistically, maresin-2 at least in part enhances cardiac autophagy by pronouncing the O-GlcNAcylation of an autophagy starter—the kinase subunit of ULK1 complex, implying a novel crosstalk between inflammation and O-GlcNAc modification.

## Methods

### Animal Preparation

The experiment protocol conformed to the Guideline for the Care and Use of Laboratory Animals published by the US National Institutes of Health (NIH Publication No. 85-23, revised 1996). All experiments were approved by the Institutional Animal Care and Use Committee of the Wuhan University and Hubei University of Arts and Science, which were also obligated to the 3R (replacement, reduction, refinement) principle. Male wild type C57BJ/6 mice were obtained from the animal experiment center of Wuhan University (no.2008-0004), China.

The global *Alox*12 knockout mice were obtained from Wuhan University to avoid the perturbation of endogenous maresin-2. The SpCas9 target was identified in the conserved exon 5 with the oligonucleotide primers annealed and cloned into *pUC57* vectors. The Cas9 nuclease encoding mRNA and *Alox12* short guide RNA (sgRNA) were introduced into C57B/J background zygotes. The phenotype of transgenic mice were examined by fast tail PCR assay kit (Beyotime, D7283M). The sequences of *Alox12* knockout were as follows: *Alox12*-sgRNA GGAGGGTATAAACACGTTTGAGG, *pUC57*-sgRNA-F GATCCCTAATACGACTCACTATAG, *pUC57*-sgRNA-R AAAAAAAGCACCGACTCGGT, *Alox12*-P1 TGGACTTTGAATGGACGTTG, *Alox12*-P2 GGGAGCACAGAAAGGACAAG.

### Animal treatment

Synthetic maresin-2 (13*R*, 14*S*-dihydroxy-4*Z*, 7*Z*, 9*E*, 11*E*, 16*Z*, 19*Z*-docosahexaenoic acid; cat# 16369, Cayman Chemical) were administered by i.p. in doses and duration set according to our experiment after referring previous study^27^. The rats were fed in a specific pathogen free environment (12/12 light-dark cycle and 65% humidity, free access to food and water). The mice underwent surgery being anesthetized with 2% sodium pentobarbital (50 mg/kg, ip), and ventilated artificially with a volume-controlled rodent respirator at 70 strokes per minute. Type I myocardial infarction (TIMI) model was established by ligating left anterior descending artery (LAD) as previously described^28^. Type II myocardial infarction (TIIMI) model was introduced by single dose of isoproterenol (100mg/kg) injection. Finishing the invasive operation, the mice were treated with 40, 000 units of penicillin. The mice were anesthetized by isoflurane inhalation when operation and sacrificed by overdose pentobarbital (250mg/kg) before sacrifice, which is a way of minimizing pain. In the process, all the operative procedures and equipment were sterile and standardized.

Auricular tragus vagus nerve stimulation (tVNS) which evidently improved cardiac vagus activity was performed as ever reported^29, 30^. Briefly, a chronic intermittent low-grade tVNS was delivered (20 Hz frequency, 1 ms pulse width, 1 mA aptitude, 1s interval bilateral) per day over the auricular cymba conchae region at each ear via a (Yuyan Ltd, Shanghai) for 30 min over 4-week period, following which myocardial infarction was induced. The electro-pulse was restrained within 1.2mV. This electric voltage is much lower than the threshold for either slowing sinus rate or decreasing epinephrine level.

### Cell preparation

Hearts were excised and washed by warm Hank’s balanced salt solution (HBSS) to remove blood before minced into the digestion reagents Liberase^TM^ TH containing collagen-I, collagen-II, and Thermolysin, a non-clostridial neural protease and DNase1 for 5min and stopped by ice cold HBSS with 10% fetal bovine serum according to previous study. Digestion finished; the triturated tissue was centrifuged. We discarded the supernatant and treated it with red blood cell lysing buffer Hybri-Max^TM^. Finally, the cells were washed twice with HBSS. NRCM were obtained from newborn mice aged 1∼2 days. HEK293 cell line was kindly provided by Hubei Key Laboratory of Cardiology. We cultured NRCM and HEK293 cell line in the DMEM/F-12 medium (SigmaAldrich, D6421) supplemented with 15mM HEPES, 0.055g/L sodium pyruvate, 3.55g/L glucose, 0.365g/L L-glutamine, 100 U/ml penicillin, and 100 ug/ml streptomycin with or without MaR2 (GLPBIO, GC40980).

### Assessment of myocardial injury

To assess the extent of myocardial injury, hematoxylin-eosin staining and sirius red staining were performed to assess the myocardial injury as previously described^29^. Blood was removed from heart samples, the sampled hearts were cut transversely into around 5 µm slices, fixed in 4% paraformaldehyde solution and embedded in paraffin for staining after deparaffinization and rehydration. The stained slices were scanned with a NanoZoomer scanner (Hamamatsu Photonics, Japan), observed by NanoZoomer Pathology Digital software NDPview2.0 and analysed via Fiji ImageJ software.

### Echocardiograph

The rats were anaesthetized with 2% barbital sodium and underwent echocardiography under unconscious conditions with a Vinno V8 ultrasound machine (Soochow, VinnoTech) and a linear array probe (4 –12 Hz) by a physician working at the imaging department of Xiangyang Central Hospital. Left ventricular ejection fraction (LVEF), left ventricular fractional shortening (LVFS), left ventricular internal diameter at end-diastole (LVIDd), and left ventricular internal diameter at end-systole (LVIDs) were measured to evaluate cardiac function and structure.

### Immunofluorescence

We fixed the animal samples in 4% paraformaldehyde before blocking them into 1% fetal bovine serum as previous study indicated. After washing, the primary antibodies including Vesicular monoamine transporter 2 (VMAT2), tyrosine hydroxylase (TH) were incubated overnight at 4^。^C. Then the samples underwent phosphate buffered saline (PBS) washing for 15min before incubated with the second antibody HRP goat anti-rabbit antibody (ThermoFisher, 31460) which was dissolved into 1% BSA for 1h in room temperature. All above finished, the second antibody was discarded by 15min PBS washing. We retained the DNA with 0.1% DAPI for incubation. After washing, the sample sheets were sealed and observed under a microscopy (Japan, Olympus).

### Transmission electron microscope (TEM)

The tissue was quickly fixed in a mixture of 4°C pre-cooling 2.5% glutaral and 0.1M phosphate purchased from Servicebio (Wuhan, G1102). The in-vitro samples were fixed in 2% glutaral at 4°C overnight. Thin cubic samples were immersed in 2% uranyl acetate and lead citrate. We detected mitochondrial condition and autophagosomes by transmission electron microscope (Japan, HITACHI HT7700 120kv) as previously described.

### Western blotting (WB) analysis and co-immunoprecipitation (coIP)

For WB the cell and tissue samples were lysed in 1x RIPA buffer and for coIP the cell lysates were prepared by IP buffer containing 50 mM Tris-HCl (pH 7.4), 10 mM NaCl, 1 mM EDTA, 0.5 mM EGTA, 1 mM MgCl2, 0.5% Triton X-100, and 1 mM phenylmethylsulphonyl fluoride as previous described. The prepared lysates for coIP need to undergo an extra centrifugation at 18,000g at 4°C for 5min before incubating with sWGA conjugated beads for 12hours. After washing with lysis buffer, the sWGA-bound protein precipitation was eluted by 2.5x sample buffer and lysis buffer for immunoblotting. Equal amounts of protein were transferred on a PVDF membrane (Merk Millipore, IPVH00010). Blocked at 5% fat-free bovine milk with PBS and 0.1% Tween-20 (Beyotime, ST825) for 1.5 h, specific primary antibodies were incubated overnight at 4°C and then incubated with secondary antibody for 1h. Before PBST (PBS+Tween-20) washing, HRP-conjugated anti-rabbit or anti-mouse HRP-conjugated anti-rabbit or anti-mouse secondary antibody were incubated for 1h. The band was developed by ChemiScope 6000.

### Measurement of the LSG neural activity

Neural activity from the LSG was recorded for 1minute. A tungsten-coated microelectrode wasinserted into the LSG, and a ground lead was con-nected to the chest wall. The electrical signalsgenerated by the LSG were recorded using a PowerLab data acquisition system (8/35, AD Instruments,Bella Vista, Australia) and amplified by an amplifer (DP-304,Warner Instruments, Hamden, Connecticut) with bandpass filters set at 3o0 Hz to 1 kHz and anamplification range of 30 to 50 times. Neural activity was defined as deflections with a signal-to-noise ratio of 3:1.

### Cytokine measurement

The main pro-inflammatory and anti-inflammatory cytokines in blood and heart were measured by corresponding ELIZA kit: tumor necrosis factor-ɑ (TNF-ɑ, Abcam, ab100747), interleukin-1β (IL-1β, Abcam, ab197742), interleukin-18 (IL-18, Abcam, ab216165), interleukin-6, (IL-6, Abcam, ab222503), interlrukin-10 (IL-10, Abcam, ab100697), transforming growth factor (TGF-β, Abcam, ab119557). G-CSF, GM-CSF, IFNγ, IL-1ɑ, IL-2, IL-4, IL-5, IL-7, IL-9, IL-12p70, IL-13, IL-17, IL-21, IL-23, MIP1ɑ, MIP1β, MIP2 were kindly provided by Hubei Key Laboratory of Cardiology. As previous done, we sealed the testing antigen in a 96-well microplate overnight at 4°C and blocked by blocking buffer. Then, we incubated the anti-rabbit IgG antibodies in the plate at room temperature for 1h before incubating HRP anti-rabbit IgG secondary antibodies. The reaction was stopped by addition of 2N H2S04 and the result of which was record at the absorbance at 450 nm on the reader.

### Metabolite detection and analysis

The metabolite analysis was carried out in BioTree Co., Ltd. Cells were washed twice with precooled physiological saline before the metabolites were extracted with cold methanol/acetonitrile (1:1, vol/vol) to remove the protein. Centrifuged for 20 minutes (14,000*g*, 4°C), the supernatant was dried in a vacuum centrifuge. For LC-MS analysis, the samples were redissolved in 100 μL acetonitrile/water (1:1, vol/vol) solvent.

### Statistical analysis

All data are expressed as mean±SD. Student t-test is used for between-group comparisons. For comparison of among groups, one-way ANOVA is used. Values were considered significantly different when P<0.05.

## Results

### 1. Maresin-2 biosynthesis was involved in cardiac sympathetic innervation

Being that preliminary data have suggested the involvement of adrenoreceptor activation in maresin-2 biosynthesis, we assessed the involvement of autonomous nervous system innervation level in maresin-2 bioavailability in heart^27^. The wild type of myocardial infarction C57BJ/6 mice was randomly separated into adrenaline induced sympathetic overdrive (SO) group and tragus vagal nerve stimulation (tVNS) group and tVNS suppressed the electric firing at a main sympathetic nodose, left stellate ganglia (Fig 1.A&B) As shown in Fig 1.C, in contrast with tVNS group, an increase of tyrosine hydroxylase (anti-TH) and vesicular monoamine transporter 2 (VMT2) were observed in SO group. To address the significance of this finding, we examined whether adding exogenous maresin-2 could in turn alter sympathetic innervation and found that maresin-2 supplement did not change the optical density of the sympathetic innervating biomarkers such as TH and VMT2 in the peri-infarction area (Fig 1.D). As shown in Fig 1.E&F, sympathetic inhibition by tVNS enhanced the expression of soluble epoxide hydroxylase (sEH) and 12-lipoxygenase (12-LOX) which synthesized maresin-2 when compared to SO group. From the data of Fig 1.G&H, maresin-2 strongly encouraged the fibrogenesis in the infarcted zone which helpful in cardiac repair, especially Col-III but spontaneously not as much prompted the aggravation of fibrosis in the border zone which deteriorated cardiac fibrosis. Because above data indicated that maresin-2 might be correlated with the autonomous nervous system (ANS), we examined whether maresin-2 was involved in the neurosecretory profiling. It turned out that the expression level of neural growth factor (NGF), brain derived neurotrophic factor (BDNF), and netrin-1 (NTN-1) were increased by maresin-2 treatment. Moreover, the profiling of inflammatory cytokines also supported the notion that maresin-2 contricuted to inhibit inflammatory cytokines, especially by repressing IL-1β, IFNγ, IL-7, IL-10, and TNFɑ(Fig 1.I). The results taken together indicated that maresin-2 was probably a spontaneous outcome of decreased sympathetic drive.

**Fig. 1.**
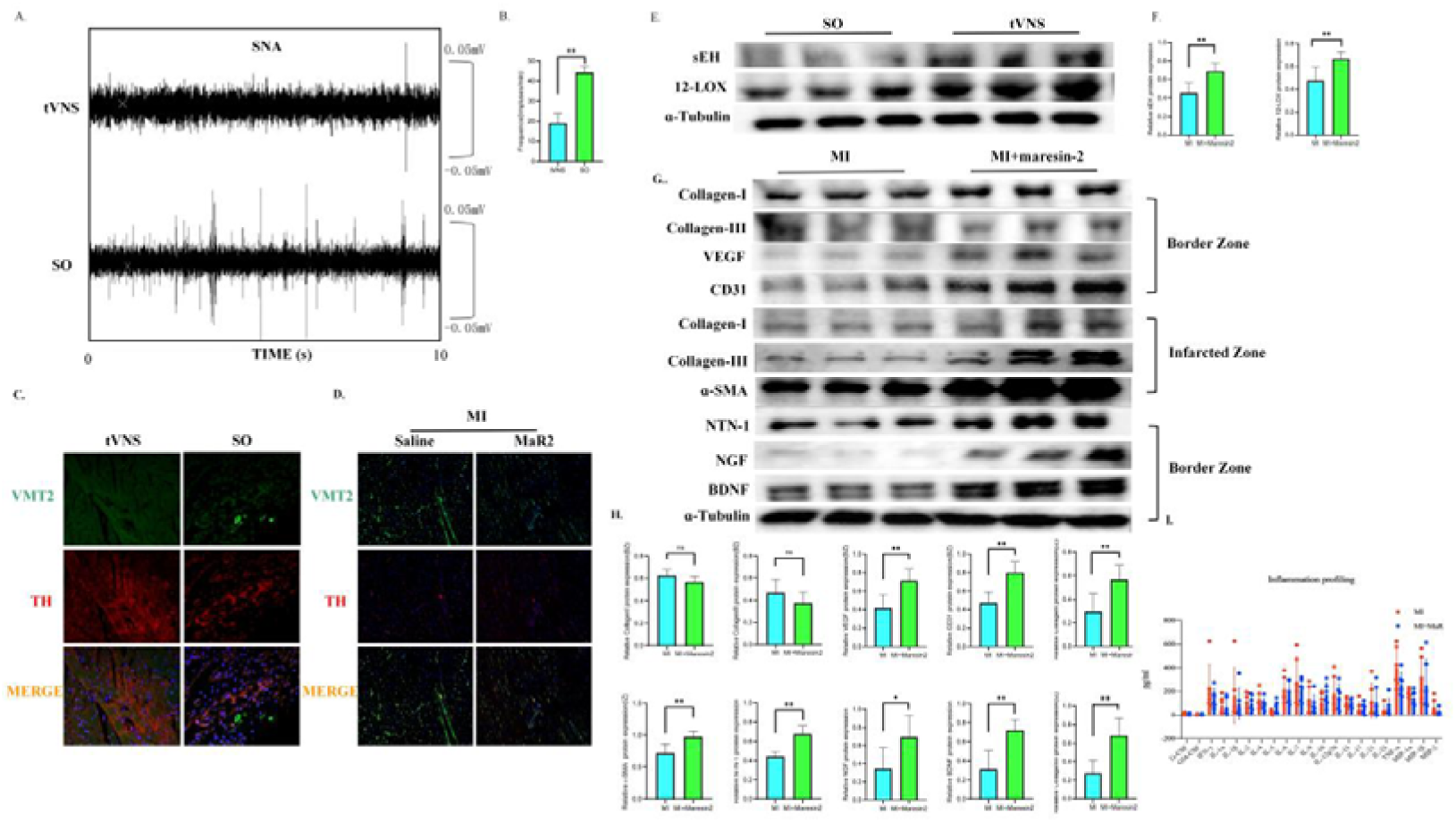
Maresin-2 biosynthesis was involved in cardiac sympathetic innervation. (A)Representative images of electrophysiology. (B) Quantitative analysis of electrophysiology. (C)Representative immunofluorescence images of GAP and TH. (D) Representative immunofluorescence images of GAP and TH. (E&F) Representative bands of western blotting of sEH and 12-LOX and quantitative analysis. (G&H) Representative bands of western blotting of ɑSMA, collagen-I, collagen-III,VEGF, CD31, NGF, BDNF, NTN-1 and quantitative analysis. (I) Major inflammation cytokines and mediators determined by Elisa. Values were means ±SEM,n=6∼9 vs MI, *P<0.05, **P<0.005.

### 2. Maresin-2 ameliorated cardiac dysfunction and limited adversary remodeling

The LAD ligated and isoproterenol injected mice aged 3∼4 weeks were fed for 2 weeks before sacrifice (Fig 2.A). As specific Alox12^-/-^ lineage mice were deprived of the potential for arachidonic metabolites biosynthesis such as maresin-1 and maresin-2 to avoid the possible bias caused by endogenous maresin-2 (Fig 2.B). We first checked the cardiac function by which we found that additional maresin-2 at baseline improved cardiac diastolic parameters of echocardiograph (Fig 2.C&D) as evident by improved left ventricular ejection fraction (LVEF), left ventricular fraction shortening (LVFS) to a detectable level but without diverging left ventricular end-systolic diameter (LVESd) and left ventricular end-diastolic diameter (LVEDd), indicating no effects on cardiac structure. However, as shown by FIHC-WGA staining (Fig 2.E) maresin-2 ameliorated the cardiac hypertrophy validated by reduced cross-sectional area per cardiomyocyte (Fig 2.F). Furthermore, results of HE and Sirius Red Staining supported that maresin-2 treated hearts were less collagen condensed in the left ventricular wall and smaller LV and RV diameters (Fig 2.G). Collectively, maresin-2 treatment improved post-infarction cardiac remodeling with improved bump function.

**Fig. 2.**
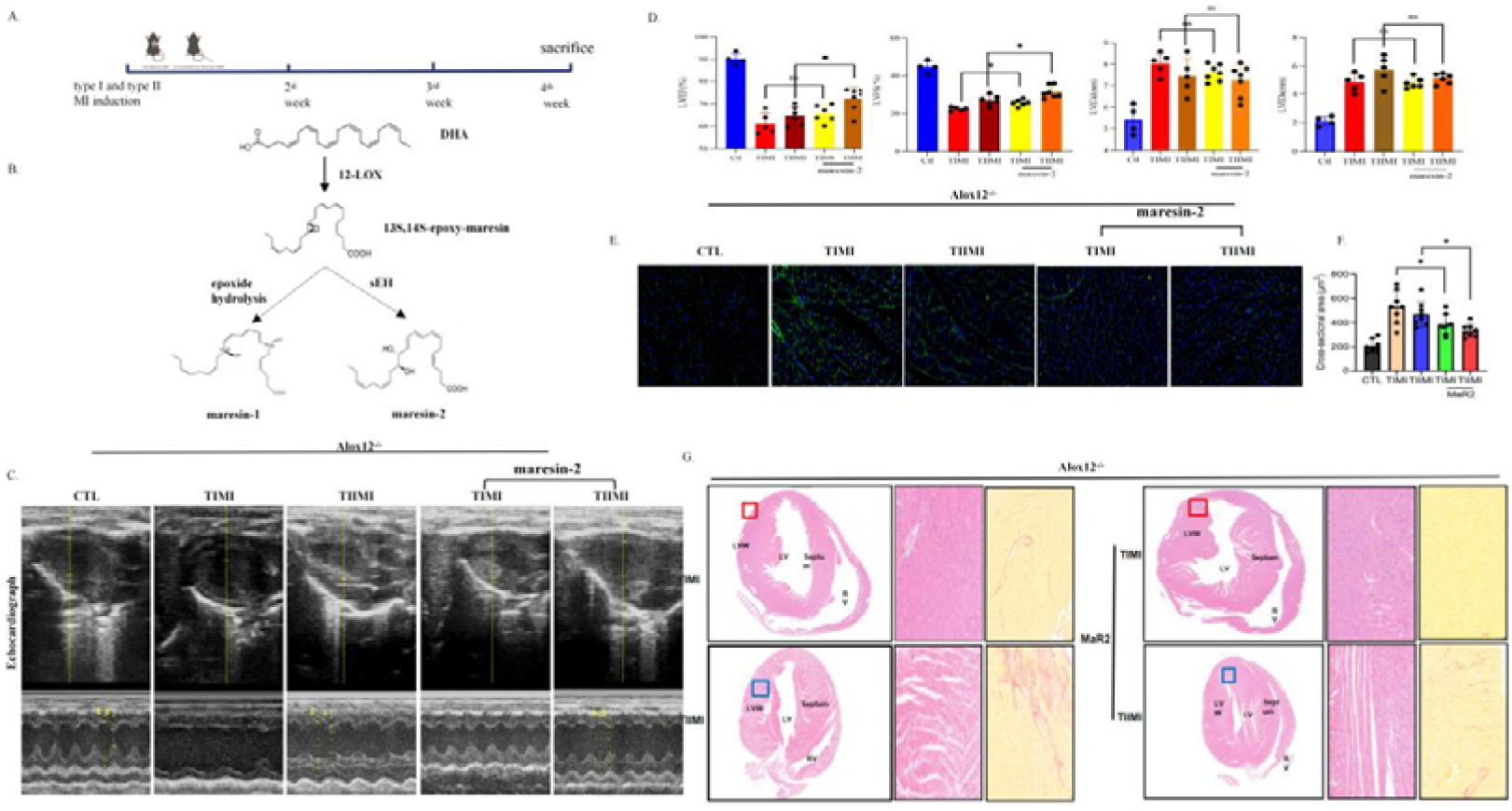
Maresin-2 ameliorated cardiac dysfunction and limited adversary remodeling. (A) The mice endergoing TIMI or TIIMI induction were sacrificed in the 4^th^ week after MI induction. (B) The schematic illustration of maresin-2 biosynthesis from polyunsaturated fatty acids such as DHA. (C) Representative images of echocardiography. (D) Quantitative analysis of cardiac function by LVEF (%), LVFS (%), LVIDd (mm), LVIDs (mm). (E) Representative images of FITC-WGA immunofluorescent staining. (F) Quantitative analysis of the cross-sectional areas of each cardiomyocytes. (G) Hematoxylin-eosin (HE) staining staining and Sirius Red staining to evaluate the myocardial injury under control and 10սg/kg/d maresin2 treatment MI mice. Values were means ±SEM, n=6∼9, vs MI, *P<0.05,**P<0.005.

### 3. Maresin-2 likely to facilitate autophagosome initiation

Unc-51 like autophagy activating kinase (ULK1) complex plays an important role in autophagosome formation, an apical autophagic process that is otherwise a category of late-phase autophagy inhibitor bafilomycin A_1_ (BafA_1_) or rapamycin, an autophagy inducer for comparison. We first of all examined the level of autophagy in primary neonatal rat ventricular cells (NRCMs) and in living mice exposed into different level of physiological concentrations of maresin-2 where we found maresin-2 dose-dependently increases the ULK1 complex expression and Bcl2 with reduced Bax (Fig 3.A&B). In the Fig 3.C&D, in fine parallel with rapamycin—the reagent bioactive to induce autophagy and against for Baf2, maresin-2 treatment also elevated the lipidated form of MAP1LC3/LC3 (microtubule associated protein 1 light chain 3) termed LC3-II while cargo receptor SQSTM1/p62 level was suppressed. It revelaed that the accumulation of autophagosomes occurs by the action of functional autophagic flux rather obstruction of autolysosome degradation. As shown in Fig 3.E, under transmission electron microscopy maresin-2 positive group exhibited more abundant autophagosome vacuoles dispatched between myofibrils, as well as less absence of normal intermyofibrillar mitochondria. We used lentivirus carrying a tandem-tagged monomeric eGFP-mCherry-LC3II reporter to transfect NRCMs (Fig. 3H&I). The steadily expressed cells were selected for assessment. Fluorescence inspection of LC3II^+^ signaling was increased in maresin-2 treatment with respect to control treatment especially in the presence hypoxia condition. Maresin-2 was not likely to interfere into the crosstalk of autophagy and apoptosis showcased by non-altered Bcl2, Bax, Caspase-3, and cleaved Caspase-3 levels in the presence or not of maresin-2 (Fig 3.F&G). However, Atg5/Atg12 dual knockdown (Atg5/Atg12 conjunct with LC3II for autophagosome nucleation) converted maresin-2 into pro-apoptotic with reduced Bcl2 and increased Bax and cleaved Caspase-3. Herein, Atg5/Atg12 complex was thought to antagonize against the potential pro-apoptosis function of maresin-2. Overall, in all likelihood, maresin-2 depends on early state autophagic process to trigger autophagosome accumulation.

**Fig. 3.**
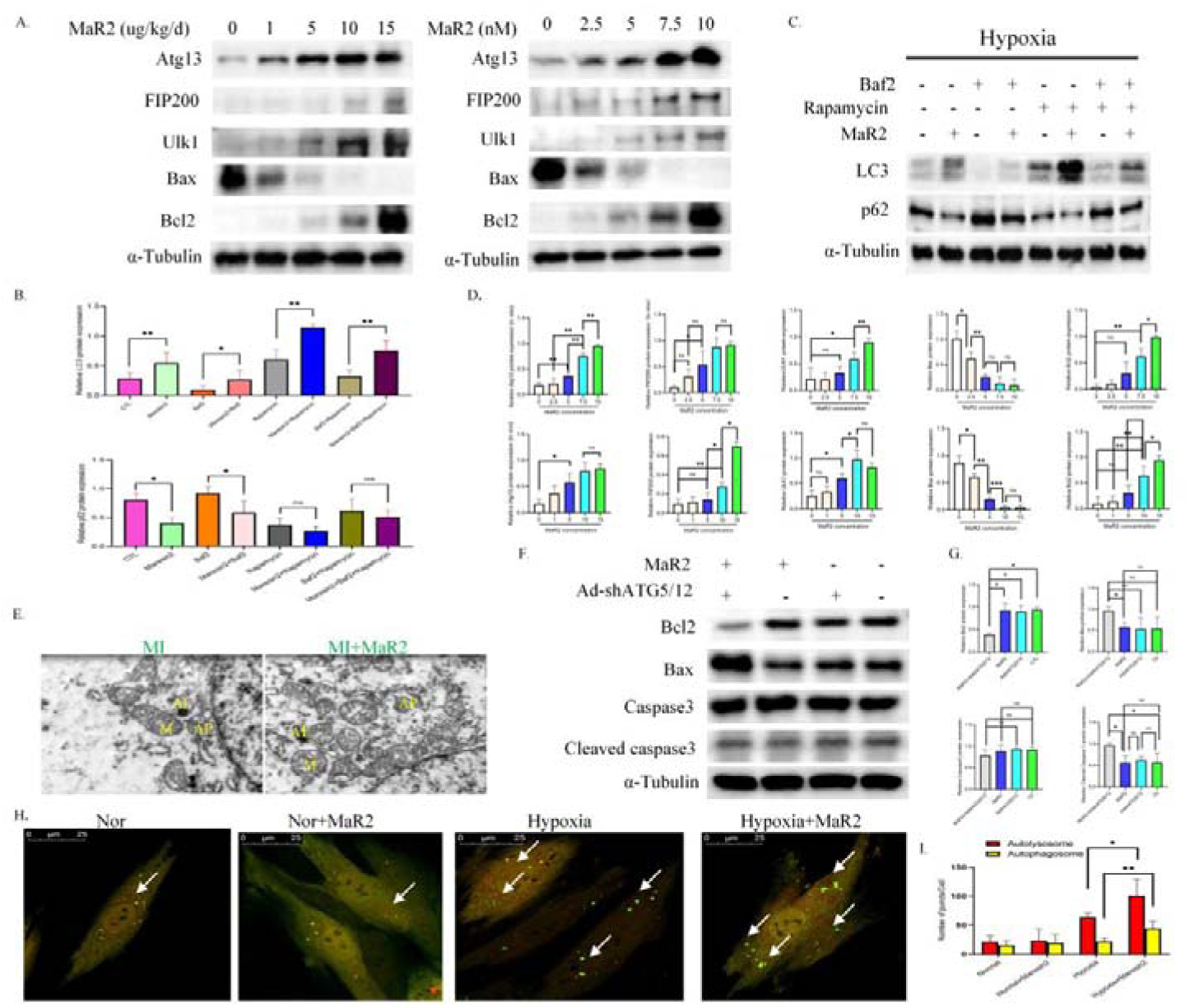
Maresin-2 likely to facilitate autophagosome initiation. (A&B) Representative western blotting bands and quantitative analysis of LC3 and p62. (C&D) Representative western blotting bands and quantitative analysis of Atg13, FIP200, Ulk1, Bax, Bcl2. (E) Representative images of autophagosome (AP), autophagolysosome (AL), and mitochondria (M) in NRCMs. (F&G) Representative western blotting bands and quantitative analysis of Bcl2, Bax, Caspase3, cleaved Caspase3 in NRCMs. (H) Autophagic flux in NRCMs was detected with the mRFP-GFP-LC3 reporters. Values were means ±SEM,n=6∼9, *P<0.05,**P<0.005.

### 4. Maresin-2 modulates ULK1 O-GlcNAcylation to induce autophagy

O-GlcNAcylation is intertwined with phosphorylation because both of them modify serine and threonine amino residues. Therefore, we assessed the O-GlcNAcylation and phosphorylation condition of ULK1. As the phosphorylation of AMPK and mTORC1, two major activators of ULK1, showed semblable oscillation amplitudes in the maresin-2 treatment group when compared with the control group (Fig 4.A&B), Supplement of Thiamet-G with Glucotamine (high O-GlcNAcylation status) strove to make a significant difference regarding autophagic flux and maintained ULK1 complex which is consistent with previous studies^32∼33^ while OSMI-1 (low O-GlcNAcylation) did exactly the same (Fig 4.C&D). Next, we tentatively assessed whether adding O-GlcNAc moiety into ULK1 the pro-autophagic mechanism of maresin-2 and harvested a significant positive result (Fig 4.E&F). In a summary, it is ULK1 O-GlcNAcylation the mechanism of maresin-2 modulated autophagy.

**Fig. 4.**
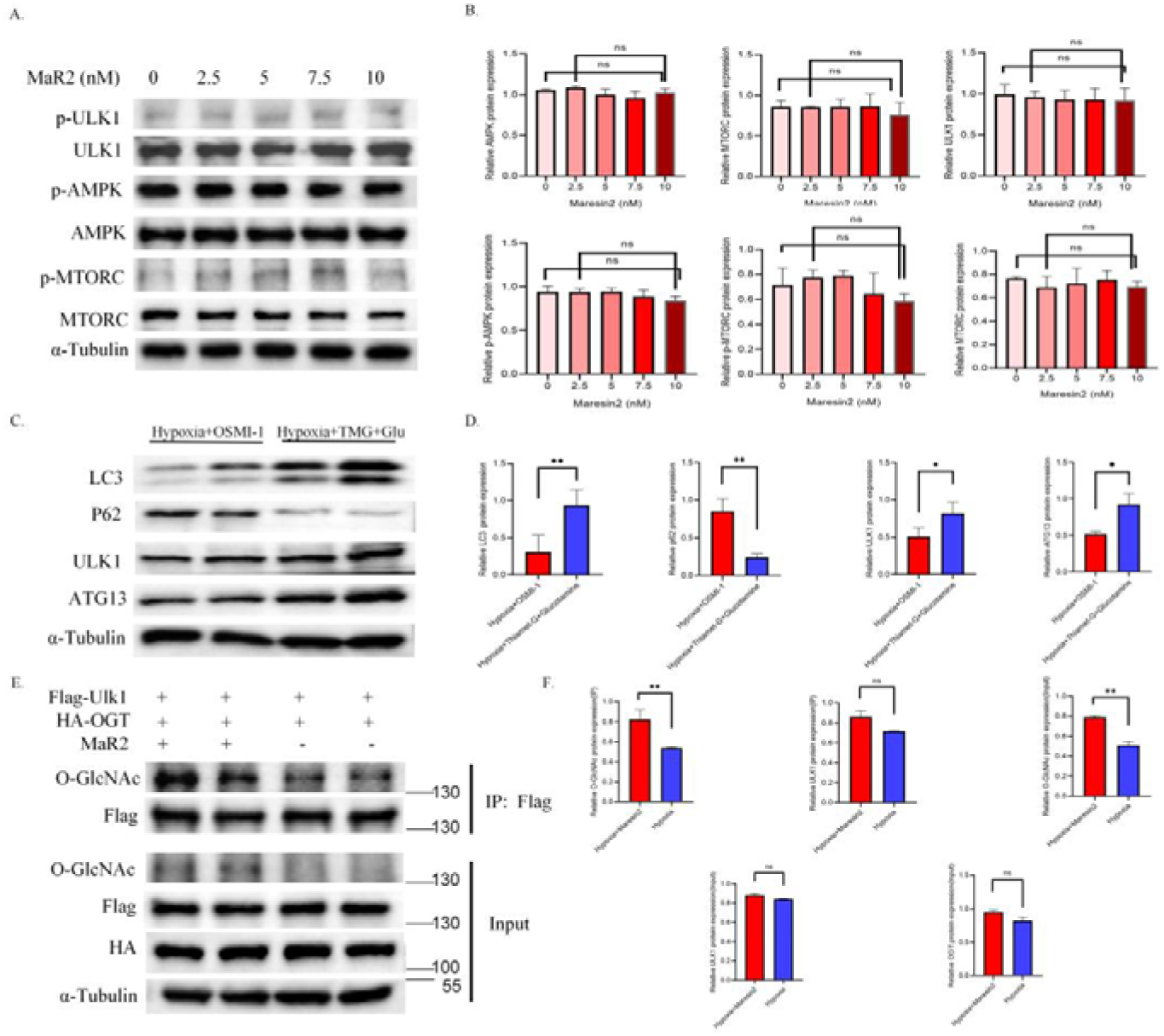
Maresin-2 modulates ULK1 O-GlcNAcylation to induce autophagy. (A&B) Representative western blotting bands and quantitative analysis of ULK1, p-ULK1, AMPK, p-AMPK, MTORC, p-MTORC under different concentrations of MaR2 in NRCMs. (C) Representative western blotting bands and quantitative analysis of LC3, p62, ULK1, ATG13 under low (OSMI-1) and high (Thiamet-G and glucosamine) O-GlcNAcylation conditions in NRCMs. (E) Immunoprecipitation assay detecting ULK1 O-GlcNAcylation under or without MaR2 supplement in HEK293 cells. Values were means ±SEM,n=6∼9, *P<0.05, **P<0.005.

### 5. Maresin-2 modifies ULK1 *O-*GlcNAcylation at Ser409/Ser422 sites

To determine the glycosylated sites of ULK1, we predicted ULK1 O-GlcNAcylation sites in an algorithm platform (http://www.cbs.dtu.dk/services/YinOYang/), which validated S298, S409, and S422 as the most vulnerable sites of O-GlcNAcylation (Fig 5.A). Ser409 is an ever-reported O-GlcNAcylated site while other fuzzy motifs never been reported. To identify the O-GlcNAc modifying sites of ULK1, we then generated Ser298Ala, Ser409Ala and Ser422Ala mutagenesis plasmids pHm*Ulk1*S298A-puroEGFP, pHm*Ulk1*S409A-puroEGFP and pHm*Ulk1*S422A-puroEGFP. The plasmids were constructed to missense mutate serine residue into alanine (Fig 5.B). Finishing the transfection of Ser409 and Ser422, maresin-2 failed to induce LC3II-mediated autophagic flux in 293T cell line with S298 site-mutation exerting no distinctive effects on autophagic flux (Fig 5.b). Above data suggested that maresin-2 possibly modified ULK1 via O-GlcNAcylation at Ser409/Ser422.

**Fig. 5.**
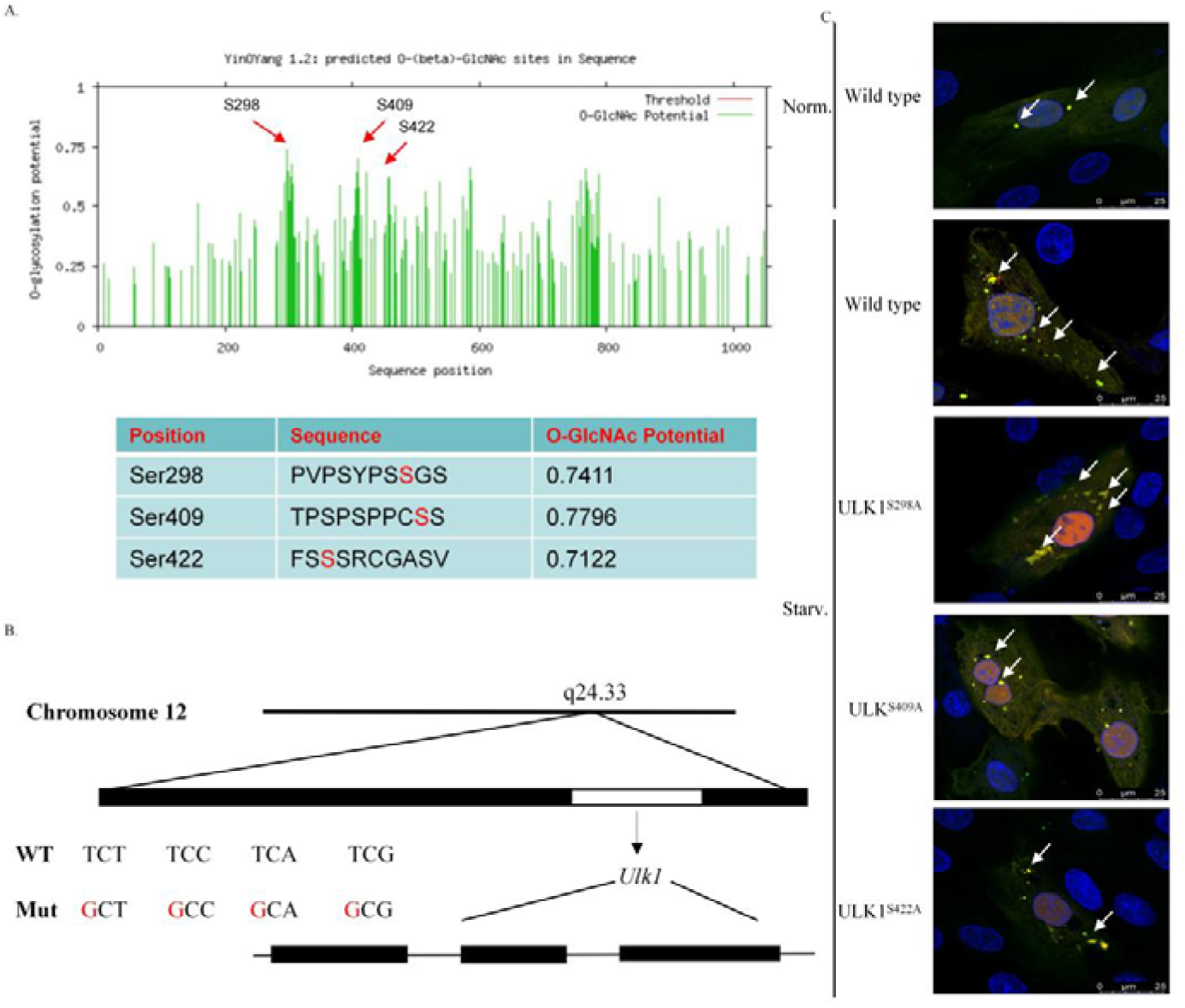
Maresin-2 modifies ULK1 O-GlcNAcylation at Ser409/Ser422 sites. (A) Three most vulnerable fuzzy O-GlcNAcylated sites of ULK1. (C) To figure out the functional potential of the predicted O-GlcNAcylation sites, we accordingly created the site-mutations on ULK1 to evaluate the implication on autophagic flux.

### 6. Maresin-2 increments hexosamine biosynthesis pathway (HBP)

Prior experiments have validated that O-GlcNAc transferase (OGT) is a highly conservative enzyme whose expression was not altered in a dynamic and reversible manner^34^. We cultured HEK293T cells with different concentration maresin-2; the GFAT1 expression level were enhanced in gradualness while the expression level of OGT, OGA, GLUT1, GLUT4 were not in statistically altered (Fig 6.A&B). Herein, we speculated that it was either glucose or glutamine fueling proteins that regulated O-GlcNAc metabolism as sketched (Fig 6.C). Importantly, UDP-GlcNAc concentration in maresin-2 present group showed twofold upsurge over blank comparation and maresin-2 also fueled the glutamine consumption which is identical with GFAT1 (Fig 6.D). Moreover, to validate the OGT/OGA activity a major factor determing maresin-2-induced O-GlcNAc modification on the ULK1, we successfully constructed a site mutation on 254 site locating at OGT tetratricopeptide repeat (TRP) domain to disrupt OGT (Fig 6.E) as proved by the decreased UDP-GlcNAc level. However, judging from the co-immunoprecipitation data (Fig 6.F), ULK1 O-GlcNAcylation was not likely to be affected by OGT^L254F^. This part of data implies that whereby hexosamine biosynthesis influx and increased rate-limiting conversion of glucosamine-6-phosphate ragther than the enzymatic activity of OGT was the very reason for maresin-2 intensified ULK1 O-GlcNAcylation.

**Fig. 6.**
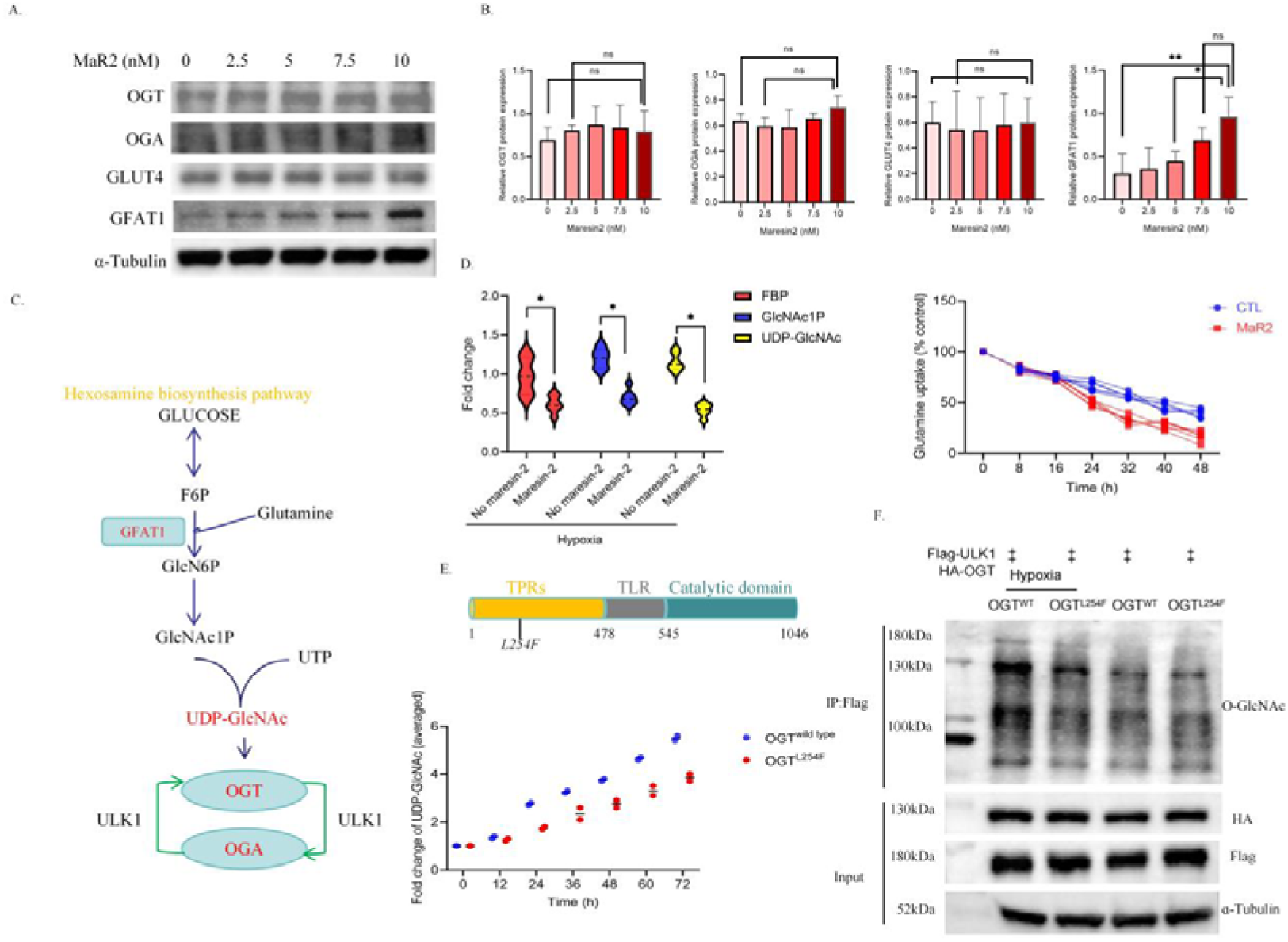
Maresin-2 increments hexosamine biosynthesis pathway (HBP) (A&B) Representative western blotting bands and quantitative analysis of OGT, OGA, GLUT1, GLUT4 under different concentrations of MaR2 in NRCMs. (C)Schematic illustration of hexosamine biosynthesis pathway which allows for protein O-GlcNAcylaton. (D) Relative UDP-GlcNAc and fructose-6-phoaphaste, and GlcNAc1P levels were measured by LC-MS in HEK293 cells. (E) To validate the effect of OGT activity on ULK1 O-GlcNAcylation, L254F-Mut-HEK293 cells were created in which the HBP was to a certain extent eliminated by a site-mutation in TRP domain. Values were means ±SEM,n=6∼9, *P<0.05, **P<0.005.

### 7. Maresin-2 renders GFAT1 activation by inhibiting TAK1-TAB1 inflammatory signalosome

Previously we considered that incremental hexosamine biosynthesis and UDP-GlcNAc amount might promote proteinic O-GlcNAcylation including ULK1^35^. D-fructose-6-phosphate aminotransferase 1 (GFAT1) which transforms glucosamine 6-phosphate into fructose 6-phosphate conjugates with TAB1 as its cofactor of hexosamine metabolism^36^. Prior literature has elucidated that TAB1 is also a cofactor of TAK1 in auxiliary responding to IL-1β, TNF-ɑ, and Toll-like receptors signaling, and as a result, NF-κB is activated. Therefore, to validate our suspicion on TAB1 activated to turn up GFAT1 activation, we scrutinized the TAB1-TAK1 inflammatory signalosome and subsequent IKKs/p65/ NF-κB activation were scrutinized (Fig 7.A&B). As shown, maresin-2 suppressed TAK1/IKKs /NF-κB phosphorylation level under the participation of TAB1. Notably, TAB1 was required for maresin-2 induced autophagy (Fig 7.C&D). Moreover, we found maresin-2 pronounced the interaction of TAB1 and GFAT1 (Fig 7.E&F) and TAB1 knockdown largely cancelled the O-terminal glycosylating modification of ULK1 (Fig 7.G&H). Altogether, above data provided a hint that maresin-2 induced ULK1 O-GlcNAcylation could be achieved via retarding TAK1-TAB1 signalosome.

**Fig. 7.**
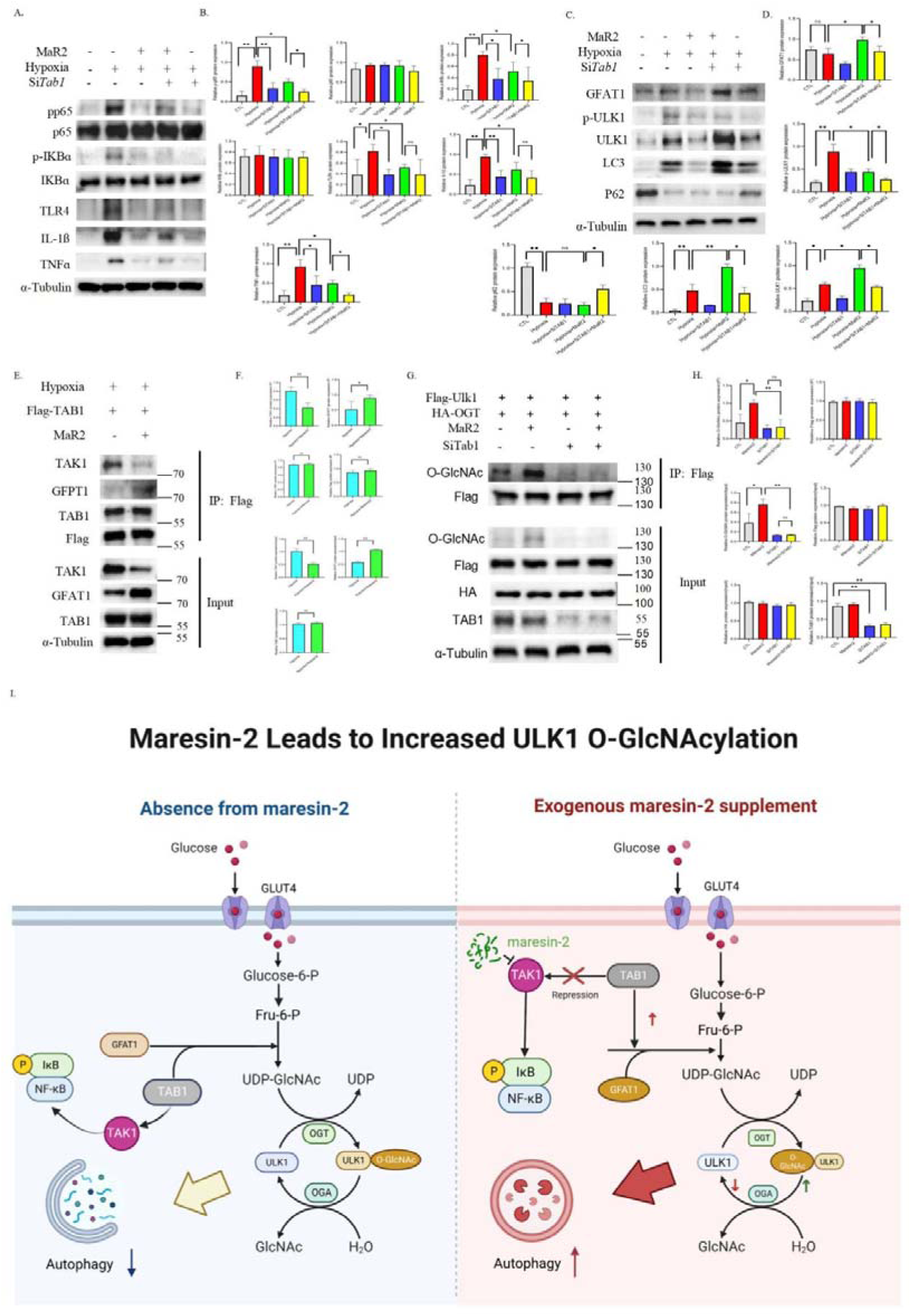
Maresin-2 renders GFAT1 activation by inhibiting TAK1-TAB1 inflammatory signalosome. Fig.7 Maresin-2 renders GFAT1 activation by inhibiting TAK1-TAB1 inflammatory signalosome (A&B)Representative bands of western blotting of pp65,p65,p-IKBɑ,IKBɑ,TLR4, IL-1ß, TNFɑ and quantitative analysis. (C&D)Representative bands of western blotting of GFPT1,p-ULK1,ULK1,LC3,P62 and quantitative analysis. (E&F) Co-immunoprecipitation of TAK1 and TAB1 or GFPT1 and TAB1 when there is MaR2 or not in HEK293 cells. (G&H)) Immunoprecipitation assay detecting ULK1 O-GlcNAcylation in HEK293 cells. I. Constructed model of our speculation. Values were means ±SEM,n=6∼9, *P<0.05, **P<0.005.

## Discussion

Here, we give hints about a previous unappreciated mechanism involving a lipokine, maresin-2 to protect against ischemic myocardial injury and attempted to demonstrate its impact on cardiac autophagy via ULK1 O-GlcNAc modification. Recent studies found endogenous maresin-2 biosynthesis promoted by environment cold is a targeted inflammatory resolution^27^. However, less is known about the role of maresin-2 on heart. Our findings suggested a possible role of maresin-2 in boosting post myocardial infarction cardiac autophagy via fine-tuning ULK1 O-GlcNAcylation. Besides, due to competitive binding of TAB1 onto either GFAT-1 or TAK1, maresin-2 treatment swings TAB1 to bind with GFAT1 for GFAT1 activation, which, subsequently leads to increased HBP influx whereby increasing O-GlcNAcylation. This “inflammation or O-GlcNAcylation” crossroad to an extent determined cardiac autophagy at least in part via ULK1 O-GlcNAcylation.

It has been well founded that polyunsaturated fatty acid are more than an energy depot; but rather a type of bioactive signaling complementary—and possibly determinant for cardiac homeostasis^37^. It was reported that disposal of DHA into maresin-1 by lipoxygenases accelerated macrophage polarizing from phagocytic milieu into regenerative phenotype and concomitantly attenuated leukocyte-mediated tissue destruction. Maresin-2 also have some tissue-reparative function although there is some functional discrepancy^38, 39^.

Autophagy is an important tissue repair machinery^40, 41^. We have explored the role of maresin-2 in programmed cell death of heart, finding that maresin-2 showcases the potential to modulate autophagy and the dynamics of ULK1 O-GlcNAcylation; Ser409 and Ser422 locating at the kinase domain of ULK1 as functional sites for glycosylating if mutatation shunts ULK1 autophagy initiation. However, our experimental setting so far cannot delineate the possible connections between the two glycosylated sites. One scenario could be Ser409 and Ser422 undergoing O-terminal glycosylation independently; or the two sites that not in the conserve domain interacted structurally to form O-GlcNAcylation. Specifically, O-GlcNAc and phosphate addition on a protein is inter-regulated either via competitively machinery conjugation or via manipulating adjacent amino acid modifications in a synergic or retardant way. The definite validation requires structural analysis such as X-ray or cryo-EM in the future.

O-GlcNAcylation has been closely linked with autophagy^42, 43^. Hexosamine biosynthesis that governs O-GlcNAcylation is highly dynamic under myocardial infarction condition and described as a double edge sword^44^. As betokened by Bin *et al.* and Pellegrini *et al*., O-GlcNAcylation predominantly involved in autophagosome-lysosome fusion by targeting the docker SNAP-29 and concomitantly participates in autophagosome membrane nucleation, as well as ATG recruitment^45, 46^. Notwithstanding, Zhang’s group dropped an opposite hint—O-GlcNAc modification of the Golgi stacking GRASSP55 hinders the interaction of LCIII and LAMP, thus suppressing autophagic flux^47^. Therefore, the effects of O-GlcNAcylation on autophagy are counteracted depending on its specification. Biochemically, ULK1, on maresin-2 treatment, showed inclined O-GlcNAcylation, likely within a considerable timeframe and not a transient thing, on two distinctive sites. Specifically, O-GlcNAc or phosphate modification is often inter-regulated either by competitively machinery conjugation or by modifying adjacent amino acid in a synergic or retardant manner. Hyper-O-GlcNAcylation (Ser409) of ULK1 is positively correlated with HSP70-mediated chaperon mediated autophagy and antagonizes against phosphorylation (Ser423)^32^. A myriad of other O-GlcNAcylation checkpoints was discussed. At first, we speculated maresin-2 altered the efficacy of OGT adding *O*-GlcNAc into substrate molecules^47^. So, we mutated some amino acid permutations in the functional OGT N-terminal tetratricopeptide repeat (TPR) domain as explored before, which consists of 13.5 repetitive 34-amino-acid antiparallel alpha helices to disrupt OGT interactome^48^. It turned out that blunted OGT interactome failed to influence ULK1 O-GlcNAcylation, indicating maresin-2 not very possibly dependent on subtle OGT activity to make a difference. Therefore, we turned to upstream checkpoints. TAB1 is not only a cofactor of NF-κB activator—TGF-β-activated kinase 1 (TAK1) but a cofactor of the HBP influx rate-limiting enzyme—GFAT1^26, 48^. GFAT1 converts fructose-6-phosphate and glutamine to glucosamine-6-phosphate precursing before UDP-GlcNAc while TAK1 activates NF-κB via IKKα***/***β, respectively^48, 49^. That is the first hypothesis, maresin-2 resolves the input of IL-1, IL-6 and TNFα signaling into heart and as a result of which, TAB1 dislocating from TAK1 turns to interact with GFAT1 to allow a pronounced HBP influx. This notion can also be supported by earlier studies that OGT is not necessarily increased upon UDP-GlcNAc accumulating, and suggest a paradox termed as “inflammation or O-GlcNAcylation” determined by with whom TAB1 is bound^50^. Another hypothesis is TAK1 suppression leads to p38 MAPK inhibition, therefore attenuating its feedback inhibition of TAB1^51^. Replenishing the TAK1-TAB1 signalosome assembly as TAB1 upregulation may seem, excessive TAB1 is sufficient to simultaneously support GFAT1 activation. This notion is in fine accordance with the fact that maresin-2 treatment negatively modulates IKβ/ NF-κB activation^52, 53^.

However, we must admit that the interplay between O-GlcNAcylation and NF-κB pathway is much more perplexed than we have discussed because NF-κB subunits themselves also undergo O-GlcNAcylation. TAB1 can be O-GlcNAcylated; it enhances the autophosphorylation of TAK1^54^. IKKβ is found to be O-GlcNAcylated; it prepares NF-κB for boosting glycolysis^55^. Our study has yet scrutinized whether maresin-2 resolves the O-GlcNAcylation of TAB-1, as well as other involved proteins such as TAK1 and IKKs in a holistic way. Moreover, to what extent O-GlcNAc modification of the NF-κB pathway subunits interfering with ULK1 O-GlcNAcylation requires further elucidation. After all, NF-κB signaling pathway is another causative factor for autophagy.

Taken together, our experiments are suggestive in finding that maresin-2 encourages O-GlcNAcylation of ULK1 to improve cardiac autophagy. These findings might serve as a springboard to understand the interaction of inflammation and O-GlcNAcylation.

## Abbreviations

LC3: microtubule-associated proteins 1A/1B light chain 3B
ULK1: unc-51 like autophagy activating kinase 1
O-GlcNAc: O-linked β-N-acetylglucosamine
GFAT1: glutamine-fructose-6-phosphate aminotransferase 2
OGT: O-linked N-acetylglucosaminyltransferase
TAK1: transforming growth factor-β (TGF-β)-activated kinase 1
TAB1: TAK1 binding protein 1
SPM: specialized prosolving mediator
DHA: docosahexaenoic acid
12-LOX: 12-lipoxygenase
ATG13: autophagy related 13 (ATG13)
ATG101: autophagy related 101
FIP200: focal adhesion kinase family interacting protein of 200 kDa
PI3P: phospholipid phosphatidylinositol 3-phosphate
VPS34 complex: vacuolar protein sorting 34 complex
OGA: O-GlcNAcase
HBP: hexosamine biosynthetic pathway
UDP-GlcNAc: uridine-5′-diphospho-*N*-acetylglucosamine
GlcN-6-P: glutamine to glucosamine-6-phosphate
NF-κB: nuclear factor kappa-light-chain-enhancer of activated B cells
p38 MAPK: p38 mitogen-activated protein kinase
sgRNA: short guide RNA
TIIMI: Type II myocardial infarction
tVNS: tragus vagus nerve stimulation
SO: sympathetic overdrive
LDH: lactate dehydrogenase
CK-MB: creatine kinase
IU: international units
CLS-IF: confocal laser scanning immunofluorescence
ɑSMA: smooth muscle actin
Col-I: collagen-I
Col-III: collagen-III
TH: tyrosine hydroxylase
VMT2: vesicular monoamine transporter 2
NGF: neural growth factor
BDNF: brain derived neurotrophic factor
NTN-1: netrin-1
LVEF: left ventricular ejection fraction,
FS: fraction shortening
ESV: end systolic volume
LVESd: end diastolic volume, left ventricular end-systolic diameter
LVEDd: left ventricular end-diastolic diameter
MTORC1: rapamycin kinase complex 1
AMPK: AMP-activated protein kinase
TPR: tetratricopeptide repeat

## Acknowledgement

Not applicable

## Ethics approval and consent to participate

This study was approved by the Institutional Animal Care and Use Committee of the Hubei University of Arts and Science, Wuhan University.

## Availability of data and material

All data and materials used during the current study are available from the corresponding author on reasonable request.

## References

1. Yap J, Irei J, Lozano-Gerona J, Vanapruks S, Bishop T, Boisvert WA. Macrophages in cardiac remodelling after myocardial infarction. Nat Rev Cardiol. 2023 Jan 10.

2. Contessotto P, Spelat R, Ferro F, Vysockas V, Krivickienė A, Jin C, Chantepie S, Chinello C, Pauza AG, Valente C, Rackauskas M, Casara A, Zigmantaitė V, Magni F, Papy-Garcia D, Karlsson NG, Ereminienė E, Pandit A, Da Costa M. Reproducing extracellular matrix adverse remodelling of non-ST myocardial infarction in a large animal model. Nat Commun. 2023 Feb 22;14(1):995.

3. Jadapalli JK, Halade GV. Unified nexus of macrophages and maresins in cardiac reparative mechanisms. FASEB J. 2018 Oct;32(10):5227–5237.

4. Li QF, Hao H, Tu WS, Guo N, Zhou XY. Maresins: anti-inflammatory pro-resolving mediators with therapeutic potential. Eur Rev Med Pharmacol Sci. 2020 Jul;24(13):7442–7453.

5. Yao D, Lv Y. A cell-free difunctional demineralized bone matrix scaffold enhances the recruitment and osteogenesis of mesenchymal stem cells by promoting inflammation resolution. Biomater Adv. 2022 Aug;139:213036.

6. Hwang SM, Chung G, Kim YH, Park CK. The Role of Maresins in Inflammatory Pain: Function of Macrophages in Wound Regeneration. Int J Mol Sci. 2019;20(23):5849.

7. López-Vicario C, Sebastián D, Casulleras M, Duran-Güell M, Flores-Costa R, Aguilar F, Lozano JJ, Zhang IW, Titos E, Kang JX, Zorzano A, Arita M, Clària J. Essential lipid autacoids rewire mitochondrial energy efficiency in metabolic dysfunction-associated fatty liver disease. Hepatology. 2022 Jul 5.

8. Schlotawa L, Lopez A, Sanchez-Elexpuru G, Tyrkalska SD, Rubinsztein DC, Fleming A. An inducible expression system for the manipulation of autophagic flux *in vivo*. Autophagy. 2022 Oct 30:1–14.

9. Ren X, Nguyen TN, Lam WK, Buffalo CZ, Lazarou M, Yokom AL, Hurley JH. Structural basis for ATG9A recruitment to the ULK1 complex in mitophagy initiation. Sci Adv. 2023 Feb 15;9(7):eadg2997.

10. Schlotawa L, Lopez A, Sanchez-Elexpuru G, Tyrkalska SD, Rubinsztein DC, Fleming A. An inducible expression system for the manipulation of autophagic flux *in vivo*. Autophagy. 2022 Oct 30:1–14.

11. Mercer CA, Kaliappan A, Dennis PB. A novel, human Atg13 binding protein, Atg101, interacts with ULK1 and is essential for macroautophagy. Autophagy. 2009 Jul;5(5):649–62.

12. Ganley IG, Lam du H, Wang J, Ding X, Chen S, Jiang X. ULK1.ATG13.FIP200 complex mediates mTOR signaling and is essential for autophagy. J Biol Chem. 2009 May 1;284(18):12297–305.

13. Egan DF, Chun MG, Vamos M, Zou H, Rong J, Miller CJ, Lou HJ, Raveendra-Panickar D, Yang CC, Sheffler DJ, Teriete P, Asara JM, Turk BE, Cosford ND, Shaw RJ. Small Molecule Inhibition of the Autophagy Kinase ULK1 and Identification of ULK1 Substrates. Mol Cell. 2015 Jul 16;59(2):285–97.

14. Stanley RE, Ragusa MJ, Hurley JH. The beginning of the end: how scaffolds nucleate autophagosome biogenesis. Trends Cell Biol. 2014 Jan;24(1):73–81.

15. Zhong Y, Tang K, Nattel S, Zhai M, Gong S, Yu Q, Zeng Y, E G, Maimaitiaili N, Wang J, Xu Y, Peng W, Li H. Myosin light-chain 4 gene-transfer attenuates atrial fibrosis while correcting autophagic flux dysregulation. Redox Biol. 2023 Apr;60:102606.

16. Zhang J, Hu S, Gao Y, Wei X, Qu Y, Gao R, Lv Y, Wang J, Wang Y, Yang J, Cao J, Zhang F, Ge J. Galangin alleviated myocardial ischemia-reperfusion injury by enhancing autophagic flux and inhibiting inflammation. Eur J Pharmacol. 2023 Feb 25:175621.

17. Ren X, Nguyen TN, Lam WK, Buffalo CZ, Lazarou M, Yokom AL, Hurley JH. Structural basis for ATG9A recruitment to the ULK1 complex in mitophagy initiation. Sci Adv. 2023 Feb 15;9(7):eadg2997.

18. Hart GW, Housley MP, Slawson C. Cycling of O-linked beta-N-acetylglucosamine on nucleocytoplasmic proteins. Nature. 2007 Apr 26;446(7139):1017–22.

19. Hanover JA, Krause MW, Love DC. Bittersweet memories: linking metabolism to epigenetics through O-GlcNAcylation. Nat Rev Mol Cell Biol. 2012 Apr 23;13(5):312–21.

20. Lv P, Du Y, He C, Peng L, Zhou X, Wan Y, Zeng M, Zhou W, Zou P, Li C, Zhang M, Dong S, Chen X. O-GlcNAcylation modulates liquid-liquid phase separation of SynGAP/PSD-95. Nat Chem. 2022 Jul;14(7):831–840.

21. Hardivillé S, Hart GW. Nutrient regulation of signaling, transcription, and cell physiology by O-GlcNAcylation. Cell Metab. 2014 Aug 5;20(2):208–13.

22. Yang X, Qian K. Protein O-GlcNAcylation: emerging mechanisms and functions. Nat Rev Mol Cell Biol. 2017 Jul;18(7):452–465.

23. Yu H, Wen L, Mu Y. O-GlcNAcylation Is Essential for Autophagy in Cardiomyocytes. Oxid Med Cell Longev. 2020 Aug 11;2020:z5602396.

24. Wang D, Hu X, Lee SH, Chen F, Jiang K, Tu Z, Liu Z, Du J, Wang L, Yin C, Liao Y, Shang H, Martin KA, Herzog RI, Young LH, Qian L, Hwa J, Xiang Y. Diabetes Exacerbates Myocardial Ischemia/Reperfusion Injury by Down-Regulation of MicroRNA and Up-Regulation of O-GlcNAcylation. JACC Basic Transl Sci. 2018 May 16;3(3):350–362.

25. Dassanayaka S, Brainard RE, Watson LJ, Long BW, Brittian KR, DeMartino AM, Aird AL, Gumpert AM, Audam TN, Kilfoil PJ, Muthusamy S, Hamid T, Prabhu SD, Jones SP. Cardiomyocyte Ogt limits ventricular dysfunction in mice following pressure overload without affecting hypertrophy. Basic Res Cardiol. 2017 May;112(3):23.

26. Wei S, Zhao Q, Zheng K, Liu P, Sha N, Li Y, Ma C, Li J, Zhuo L, Liu G, Liang W, Jiang Y, Chen T, Zhong N. GFAT1-linked TAB1 glutamylation sustains p38 MAPK activation and promotes lung cancer cell survival under glucose starvation. Cell Discov. 2022 Aug 9;8(1):77.

27. Sugimoto S, Mena HA, Sansbury BE, Kobayashi S, Tsuji T, Wang CH, Yin X, Huang TL, Kusuyama J, Kodani SD, Darcy J, Profeta G, Pereira N, Tanzi RE, Zhang C, Serwold T, Kokkotou E, Goodyear LJ, Cypess AM, Leiria LO, Spite M, Tseng YH. Brown adipose tissue-derived MaR2 contributes to cold-induced resolution of inflammation. Nat Metab. 2022 Jun;4(6):775–790.

28. Snider, J.C., Riley, L.A., Mallory, N.T., Bersi, M.R., Umbarkar, P., Gautam, R., Zhang, Q., Mahadevan-Jansen, A., Hatzopoulos, A.K., Maroteaux, L., Lal, H., Merryman, W.D.,2021. Targeting 5-HT2B receptor signaling prevents border zone expansion and improves microstructural remodeling after myocardial infarction. Circulation 143,1317–1330.

29. Yang Y, Xie L, Peng Y, Yan H, Huang J, Xiao Z, Lu X. Single-Cell Transcriptional Profiling Reveals Low-Level Tragus Stimulation Improves Sepsis-Induced Myocardial Dysfunction by Promoting M2 Macrophage Polarization. Oxid Med Cell Longev. 2022 Oct 15;2022:3327583.

30. Lai Y, Zhou X, Guo F, Jin X, Meng G, Zhou L, Chen H, Liu Z, Yu L, Jiang H. Non-invasive transcutaneous vagal nerve stimulation improves myocardial performance in doxorubicin-induced cardiotoxicity. Cardiovasc Res. 2022 Jun 22;118(7):1821–1834.

31. Song J, Liu J, Cui C, et al. Mesenchymal stromal cells ameliorate diabetes-induced muscle atrophy through exosomes by enhancing AMPK/ULK1-mediated autophagy [published online ahead of print, 2023 Jan 27]. J Cachexia Sarcopenia Muscle. 2023;10.1002/jcsm.13177.

32. Shi Y, Yan S, Shao GC, et al. O-GlcNAcylation stabilizes the autophagy-initiating kinase ULK1 by inhibiting chaperone-mediated autophagy upon HPV infection. J Biol Chem. 2022;298(9):102341.

33. Yang Y, Yan Y, Yin J, et al. O-GlcNAcylation of YTHDF2 promotes HBV-related hepatocellular carcinoma progression in an N^6^-methyladenosine-dependent manner. Signal Transduct Target Ther. 2023;8(1):63.

34. Li X, Lei C, Song Q, et al. Chemoproteomic profiling of O-GlcNAcylated proteins and identification of O-GlcNAc transferases in rice. Plant Biotechnol J. 2022;10.1111/pbi.13991.

35. Wu J, Tan Z, Li H, et al. Melatonin reduces proliferation and promotes apoptosis of bladder cancer cells by suppressing O-GlcNAcylation of cyclin-dependent-like kinase 5. J Pineal Res. 2021;71(3): e12765.

36. Wei S, Zhao Q, Zheng K, et al. GFAT1-linked TAB1 glutamylation sustains p38 MAPK activation and promotes lung cancer cell survival under glucose starvation. Cell Discov. 2022;8(1):77.

37. Wu J, Liang J, Huang Y. Circulating Vegetable Omega-3 and Prognosis in Patients With Heart Failure: More Data Are Needed. J Am Coll Cardiol. 2023;81(9):e67.

38. Yu CX, Shi ZA, Ou GC, et al. Maresin-2 alleviates allergic airway inflammation in mice by inhibiting the activation of NLRP3 inflammasome, Th2 type immune response and oxidative stress. Mol Immunol. 2022;146:78–86.

39. Olsen MV, Lyngstadaas AV, Bair JA, et al. Signaling Pathways Used by the Specialized Pro-Resolving Mediator Maresin 2 Regulate Goblet Cell Function: Comparison with Maresin 1. Int J Mol Sci. 2022;23(11):6233.

40. Wu T, Zhang X, Liu Y, Cui C, Sun Y, Liu W. Wet adhesive hydrogel cardiac patch loaded with anti-oxidative, autophagy-regulating molecule capsules and MSCs for restoring infarcted myocardium. Bioact Mater. 2022;21:20–31.

41. Park JH, Kim H, Moon HR, et al. Human cardiac stem cells rejuvenated by modulating autophagy with MHY-1685 enhance the therapeutic potential for cardiac repair. Exp Mol Med. 2021;53(9):1423–1436.

42. Yokoe S, Hayashi T, Nakagawa T, et al. Augmented O-GlcNAcylation exacerbates right ventricular dysfunction and remodeling via enhancement of hypertrophy, mitophagy, and fibrosis in mice exposed to long-term intermittent hypoxia. Hypertens Res. 2023;46(3):667–678.

43. Luo R, Li G, Zhang W, et al. O-GlcNAc transferase regulates intervertebral disc degeneration by targeting FAM134B-mediated ER-phagy. Exp Mol Med. 2022;54(9):1472–1485.

44. Li Q, Taegtmeyer H, Wang ZV. Diverging consequences of hexosamine biosynthesis in cardiovascular disease. J Mol Cell Cardiol. 2021;153:104–105.

45. Guo B, Liang Q, Li L, et al. O-GlcNAc-modification of SNAP-29 regulates autophagosome maturation. Nat Cell Biol. 2014;16(12):1215–1226.

46. Pellegrini FR, De Martino S, Fianco G, et al. Blockage of autophagosome-lysosome fusion through SNAP29 O-GlcNAcylation promotes apoptosis via ROS production. Autophagy. 2023;1–16.

47. Zhang X, Wang L, Lak B, Li J, Jokitalo E, Wang Y. GRASP55 Senses Glucose Deprivation through O-GlcNAcylation to Promote Autophagosome-Lysosome Fusion. Dev Cell. 2018;45(2):245–261.e6.

48. Guo G, Gong K, Beckley N, et al. EGFR ligand shifts the role of EGFR from oncogene to tumour suppressor in EGFR-amplified glioblastoma by suppressing invasion through BIN3 upregulation. Nat Cell Biol. 2022;24(8):1291–1305.

49. Kato N, Dasgupta R, Smartt CT, Christensen BM. Glucosamine:fructose-6-phosphate aminotransferase: gene characterization, chitin biosynthesis and peritrophic matrix formation in Aedes aegypti. Insect Mol Biol. 2002;11(3):207–216.

50. Hu J, Gao Q, Yang Y, et al. Hexosamine biosynthetic pathway promotes the antiviral activity of SAMHD1 by enhancing O-GlcNAc transferase-mediated protein O-GlcNAcylation. Theranostics. 2021;11(2):805–823.

51. Chang MC, Chen NY, Chen JH, et al. bFGF stimulated plasminogen activation factors, but inhibited alkaline phosphatase and SPARC in stem cells from apical Papilla: Involvement of MEK/ERK, TAK1 and p38 signaling. J Adv Res. 2022;40:95–107.

52. Yu CX, Shi ZA, Ou GC, et al. Maresin-2 alleviates allergic airway inflammation in mice by inhibiting the activation of NLRP3 inflammasome, Th2 type immune response and oxidative stress. Mol Immunol. 2022;146:78–86.

53. Jadapalli JK, Halade GV. Unified nexus of macrophages and maresins in cardiac reparative mechanisms. FASEB J. 2018;32(10):5227–5237.

54. Papanicolaou KN, Jung J, Ashok D, et al. Inhibiting O-GlcNAcylation impacts p38 and Erk1/2 signaling and perturbs cardiomyocyte hypertrophy. J Biol Chem. 2023;299(3):102907.

55. Pathak S, Borodkin VS, Albarbarawi O, Campbell DG, Ibrahim A, van Aalten DM. O-GlcNAcylation of TAB1 modulates TAK1-mediated cytokine release. EMBO J. 2012;31(6):1394–1404.

